# Proteomic analysis reveals APOE isoform-specific regulation of ribosomes in neural precursor cells

**DOI:** 10.1101/2023.07.15.549177

**Authors:** M Srividya, Pradip Paul, Sarayu Ramakrishna, Vivek Ghose, Gourav Dey, Ravi S Muddashetty, Sanjeev Jain, Meera Purushottam, Biju Viswanath, Reeteka Sud

**Affiliations:** NIMHANS Bangalore; IISc Bangalore; Institute of Bioinformatics, Bangalore; Nimhans; National Institute of Mental Health and Neuro Sciences

## Abstract

ApoE4 isoform contributes to increased risk for Alzheimer’s Disease (AD) over the life course of individuals. Much remains unknown about the biological pathways that connect *APOE4* genotype with the development of pathology that eventually leads to AD, nor do we know how early in life these cellular alterations begin. To answer these questions, we derived neural precursor cells (NPCs) from induced pluripotent stem cells (IPSCs) that were CRISPR-edited at the *APOE* locus. We intended to characterize the protein expression landscape in the NPCs subsequent to targeted deletion of E4 from a parent IPSC line of *APOE3/4* genotype. Differentially expressed proteins (DEPs) following mass spectrometric analysis were determined from the protein abundance fold change values obtained for each protein. Proteins which showed >1.5-fold difference with FDR adjusted P-value < 0.05 were considered differentially expressed. DEPs were mapped to the STRING database (v11.5) for retrieval of interacting proteins and functional enrichment. CRISPR-editing of E4 from the parent line revealed 98 differential expressed proteins. Of these, 54 were upregulated, and 44 were downregulated. Further analysis of the DEPs via STRING database showed that these changes primarily affect pathways linked to RNA processing, plasma membrane repair, and cytoskeleton organization. Indeed, we find the effects of E4 extend beyond proteins considered central to AD pathology. Knowing more about the protein interactions regulated by ApoE, in an isoform-specific manner, can reveal new mechanistic insights into development of AD.

## Introduction

APOE is well known and extensively investigated for the isoform-specific increased risk it confers towards Alzheimer’s disease (AD). Though known predominantly for its role in lipid transport, the encoded protein (ApoE) also modulates various intracellular processes, including cytoskeletal assembly and stability, mitochondrial integrity and function, dendritic morphology and function ([Huang and Mahley., 2014](https://www.ncbi.nlm.nih.gov/pmc/articles/PMC4253862/#:~:text=A%20major%20function%20of%20apoE,to%20specific%20cell%2Dsurface%20receptors)).

In the CNS, APOE is primarily expressed by astrocytes and to a lesser extent by pericytes, oligodendrocytes, choroid plexus, and neurons under stressed conditions ([Achariyar et al., 2016](https://pubmed.ncbi.nlm.nih.gov/27931262/); [Bruinsma et al., 2010](https://www.sciencedirect.com/science/article/abs/pii/S0006899309027085?via%3Dihub); [Nelissen et al., 2012](https://onlinelibrary.wiley.com/doi/abs/10.1002/jnr.22743); [Pitas et al., 1987](https://www.sciencedirect.com/science/article/abs/pii/0005276087902955); [Xu et al., 2006](https://pubmed.ncbi.nlm.nih.gov/16687490/)). It is released into the CSF bound to lipids (hence called the lipidated version) by glial cells. The three different isoforms, ApoE2, ApoE3, and ApoE4 have different levels of lipidation ([Hu et al. 2015](https://pubmed.ncbi.nlm.nih.gov/25871773/)). Individuals that express E4 have lower levels of lipidated ApoE compared to those who have E3. Beyond its principal function, ApoE isoforms also have functional consequences for cognition and disease onset in AD. Not surprising then, that APOE genotype is also tightly connected to brain structure. Interestingly, this influence could begin during neurodevelopmental period: infants, as young as 2-6 months, with APOE4 allele show slower rate of white matter myelin and gray matter volume ([Dean et al. 2014](https://www.ncbi.nlm.nih.gov/pmc/articles/PMC4056558/)). More recent studies have documented altered cognitive development associated with these structural changes ([Remer et al. 2020](https://www.ncbi.nlm.nih.gov/pmc/articles/PMC7779366/)).

However, the biological processes through which ApoE modulates the early development of the brain remain unclear. It is crucial to understand the developmental roles of ApoE and the isoform-specific differences in its functions to better understand its function and dysfunction. To this end, we used neural precursor cells (NPCs) derived from induced pluripotent stem cells (iPSCs) that are CRISPR-edited at the *APOE* locus for the present study ([Schmid et al., 2021](https://www.sciencedirect.com/science/article/pii/S187350612100026X)). Proteomic analysis of these NPCs show that targeted removal of E4 differential expression of proteins involved in rRNA processing and ribosome biogenesis, and fatty acid and lipid metabolism.

## METHODS

### Induced pluripotent stem cell (iPSC) maintenance

The iPSCs [APOE 2/-, APOE 3/-, APOE 4/-, and APOE 3/4] were obtained from Bioneer A/S. Briefly, the iPSCs from an 18-year-old male of APOE 3/4 genotype were subjected to clustered regularly interspaced short palindromic repeats (CRISPR)-Caspase 9 (Cas9) gene editing to obtain isogenic iPSC lines of APOE 2/-, APOE 3/- and APOE 4/- genotype ([Schmid et al., 2019](https://pubmed.ncbi.nlm.nih.gov/30660866/)). The iPSCs were maintained in mTeSR1 complete media (72 232, Stem Cell Technologies) at 37°C, 5% CO2 conditions. A mixture of 1-mg/ml Collagenase IV (17104019, ThermoFisher Scientific), 0.25% Trypsin, 20% KO serum (10828028, ThermoFisher Scientific), 1 mm calcium chloride (made in PBS) was used to dissociate the iPSCs for passaging.

### Generation of NPCs from human iPSCs

iPSCs were characterized for pluripotency markers OCT4 and NANOG by immunostaining. Using StemPro Accutase (Gibco), the iPSC culture was enzymatically dissociated, then grown in suspension until day 7 in Embryoid Body (EB) medium. [Knockout DMEM (Gibco), 20% KOSR (Gibco), 0.1 mM Non-Essential Amino Acids (Gibco), 2 mM Glutamax, 1% Penicillin-Streptomycin (Gibco), and 0.1 mM Beta-Mercaptoethanol (Gibco)]. From day 7 to day 14, EB medium was replaced for Neural Induction Medium. [DMEM/F12 (Gibco), N2 supplement (Gibco), 8 ng/ml bFGF (Gibco), 1x Glutamax (Gibco), 1x Penicillin-Streptomycin (Gibco), 1x Non-essential Amino Acids (Gibco) and 2 µg/ml Heparin (Sigma)]. The EBs were plated on dishes coated with Matrigel (Corning) and allowed to develop neural rosettes. After manually passaging the neural rosettes, the tertiary rosettes were mechanically dissociated via pipetting and plated as an NPC monolayer. The medium was then replaced with Neural Expansion Medium [DMEM/F12 (Gibco), N2 supplement (Gibco), B27 supplement without Vitamin A (Gibco), 8 ng/ml bFGF (Gibco), 1xGlutamax (Gibco), 1x Penicillin-Streptomycin (Gibco), 1x Non-essential Amino Acids (Gibco) and 2 µg/ml Heparin (Sigma)]. Quantitative immunolabeling with Nestin and Pax6 revealed comparable NPC differentiation from iPSC lines. Cellular characterization of NPCs was performed by immunocytochemistry for Pax6 and Sox2.

### Mass spectrometry and proteomic analysis

Protein isolates from NPCs were analyzed on the Orbitrap Fusion Tribrid mass spectrometer (Thermo Fisher Scientific, Bremen, Germany). MS data analysis was carried out by using the Proteome Discoverer platform, version 2.2 (Thermo Scientific). The data were searched against NCBI Human RefSeq 89 database, which contained 110,386 unique protein sequences with known contaminants using SequestHT (Version 2.4) search algorithms. The search parameters used were set as indicated—precursor mass tolerance was set to 10 ppm and fragment mass tolerance to 0.05 Da. Oxidation of methionine and acetylation at protein N-terminus was set as variable modification, while carbamidomethylation of cysteine was set as a static modification. Other search parameters included two missed cleavages by trypsin and a 1% false discovery rate (FDR).

### Differential protein expression analysis

Proteins that showed >1.5-fold difference with FDR adjusted q-value <0.05 were considered differentially expressed (DEPs). The DEPs were also input into The Database for Annotation, Visualization and Integrated Discovery (DAVID) 2021 to identify the enriched functional pathways ([Huang et al., 2009](https://pubmed.ncbi.nlm.nih.gov/19131956/); [Sherman et al., 2022](https://pubmed.ncbi.nlm.nih.gov/35325185/)).

### Immunostaining and nucleoli quantification

The NPCs grown on coverslips were subjected to immunostaining of the nucleolar protein Fibrillarin. The cells were fixed with 4% PFA for 1 minute, followed by ice-cold methanol for 1 minute, which was followed by three washes with 1× PBS. This was followed by permeabilization with 0.1% Triton X-100 made in 1× PBS. This was followed by 1-h blocking with 3% BSA prepared in 1× PBS. They were incubated with the primary antibody (prepared in 1% BSA blocking buffer) (Fibrillarin - Abcam Cat#AB4566) overnight at 4°C. This was followed by three washes with 1× PBS and 1-h incubation with the secondary antibody (prepared in 1× PBS) at room temperature. after three washes with 1× PBS, the coverslips were mounted with Vectashield Vibrance with DAPI mounting media (Vector Laboratories - Cat#H-1800-2). The coverslips were imaged on Zeiss inverted fluorescence microscope with 63× objective. The number of nucleoli per nucleus was quantified using a CellProfiler (Carpenter et al., 2006) pipeline.

## RESULTS

### Proteomic Analysis

Neural Precursor Cells (NPCs) were derived from commercially available iPSC lines with the genotypes *APOE 3/4* (parental) and *APOE 3/-* (CRISPR-edited). These NPCs were used to perform mass spectrometry, followed by proteomic analysis.

Proteins that showed >1.5-fold difference with FDR adjusted q-value <0.05 were considered differentially expressed (DEPs). CRISPR-mediated removal of APOE4 from the parent line revealed 98 DEPs, of which 54 were upregulated, and 44 were downregulated (Table 1). Further analysis of the DEPs via the STRING database showed that these changes primarily affect pathways linked to RNA processing, plasma membrane repair, and cytoskeleton organization.

**Table 1:** Fold changes values for each of the 98 proteins that were differentially expressed in APOE3/- compared to the parental line. FC values in red shows upregulated DEPs, and blue shows downregulated DEPs.

| APOE 3/- vs 3/4 |  | APOE 3/- vs 3/4 |  |
| --- | --- | --- | --- |
| Protein | Fold Change | Protein | Fold Change |
| ADARB2 | 1.596 | ANXA3 | 0.55 |
| ALB | 9.106 | ANXA6 | 0.579 |
| ALDH1A1 | 3.756 | BASP1 | 0.487 |
| ARHGAP19 | 1.639 | BCAM | 0.627 |
| B3GAT3 | 1.857 | CALD1 | 0.535 |
| BHMT | 1.932 | CALD1 | 0.513 |
| CCZ1 | 2.225 | CAST | 0.639 |
| CIAO2B | 1.801 | CPNE4 | 0.432 |
| CLPX | 1.629 | CSRP1 | 0.663 |
| ENPP4 | 1.648 | CTSA | 0.654 |
| FADS2 | 1.843 | ELAVL4 | 0.615 |
| FGD2 | 7.513 | FZD5 | 0.36 |
| FKBPL | 1.612 | HIGD2A | 0.552 |
| FPGS | 2.036 | HIST1H1D | 0.3 |
| GPAA1 | 3.335 | HIST2H3PS2 | 0.342 |
| IDE | 1.814 | HMGB3 | 0.571 |
| INPP4A | 1.561 | HMGN1 | 0.375 |
| IPO4 | 1.583 | ING4 | 0.254 |
| KDM4D | 2.676 | ITGB1 | 0.666 |
| LGALS8 | 2.143 | MANF | 0.622 |
| LIPG | 2.272 | MPZL1 | 0.633 |
| LOC105371727 | 1.542 | MSMO1 | 0.36 |
| LPCAT1 | 1.606 | MXRA7 | 0.657 |
| MAGT1 | 1.615 | MYEF2 | 0.559 |
| MRM1 | 4 | NABP2 | 0.376 |
| NCOA5 | 2.457 | PABPN1 | 0.661 |
| NDEL1 | 5.345 | PRCP | 0.556 |
| NDN | 1.538 | PRKAG1 | 0.662 |
| NDUFB4 | 1.585 | PTGR1 | 0.584 |
| NEFL | 1.62 | SH3BGRL | 0.588 |
| NKIRAS2 | 1.931 | SIX3 | 0.573 |
| NOL11 | 1.872 | SLC39A6 | 0.523 |
| NRM | 1.71 | SRSF4 | 0.624 |
| PLCD3 | 2.438 | STMN1 | 0.632 |
| POLR3K | 1.693 | TAGLN | 0.52 |
| PUS3 | 3.734 | TAX1BP3 | 0.563 |
| RCL1 | 6.425 | TCEAL3 | 0.358 |
| RPL22 | 1.636 | TMSB15B | 0.471 |
| RPP38 | 1.906 | TMSB4X | 0.423 |
| SCD | 3.57 | TPM1 | 0.559 |
| SMYD5 | 1.521 | TPM1 | 0.526 |
| SNX15 | 1.686 | TPPP3 | 0.565 |
| TCAF1 | 1.664 | TRAPPC1 | 0.563 |
| TCF3 | 1.619 | TSPAN6 | 0.508 |
| TMEM38B | 1.826 |  |  |
| TPBG | 2.039 |  |  |
| TPD52L1 | 1.781 |  |  |
| TRIM26 | 1.783 |  |  |
| TRMT112 | 1.653 |  |  |
| TRPC4AP | 2.191 |  |  |
| TXNDC12 | 1.521 |  |  |
| UTP25 | 1.914 |  |  |
| WLS | 3.81 |  |  |
| XAB2 | 1.548 |  |  |

**Table 1:** Fold changes values the proteins that were differentially expressed in APOE2/- and APOE4/- compared to the parental line. FC values in red shows upregulated DEPs, and blue shows downregulated DEPs.

| APOE 2/- vs 3/4 |  |
| --- | --- |
| Protein | Fold Change |
| RCL1 | 3.79 |
| PUS3 | 2.081 |
| TNRC6C | 1.627 |
| ANXA3 | 0.561 |
| CALD1 | 0.524 |
| CCDC92B | 0.361 |

| APOE 4/- vs 3/4 |  |
| --- | --- |
| Protein | Fold Change |
| NEFL | 1.996 |
| ANXA3 | 0.66 |

The online database DAVID was used to identify the biological processes enriched by the DEPs. The analysis showed significant enrichment for the following processes: rRNA processing, ribosome biogenesis, lipid biosynthesis, lipid metabolism, Fatty acid biosynthesis, and Fatty acid metabolism. The DEPs enriched for ribosomal biogenesis and rRNA processing include IPO4, MRM1, NOL11, RPP38, TRMT112, and RCL1. The DEPs enriched for fatty acid and lipid metabolic processes include ALDH1A1, FADS2, INPP4A, LIPG, LPCAT1, MSMO1, PLCD3, PTGR1,

PRKAG1, and SCD (Table 3).

**Table 3:** List of DEPs enriched for biological pathways using DAVID Analysis. FC values in red shows upregulated DEPs, and blue shows downregulated DEPs.

| Lipid/Fatty Acid Metabolism |  |
| --- | --- |
| Protein | Fold Change |
| ALDH1A1 | 3.756 |
| FADS2 | 1.843 |
| INPP4A | 1.561 |
| LIPG | 2.272 |
| LPCAT1 | 1.606 |
| PLCD3 | 2.438 |
| SCD | 3.57 |
| MSMO1 | 0.36 |
| PRKAG1 | 0.662 |
| PTGR1 | 0.584 |

| Ribosomal Biogenesis |  |
| --- | --- |
| Protein | Fold Change |
| IPO4 | 1.583 |
| MRM1 | 4 |
| NOL11 | 1.872 |
| RCL1 | 6.425 |
| RPP38 | 1.906 |
| TRMT112 | 1.653 |

**Table 4:** Fold changes values the proteins that were differentially expressed in APOE2/- and APOE4/- compared to the APOE3/- (FC values in red shows upregulated DEPs, and blue shows downregulated DEPs).

| APOE 4/- vs 3/- |  |
| --- | --- |
| Protein | Fold Change |
| HMGN1 | 2.557 |
| TCEAL3 | 2.35 |
| MSMO1 | 2.286 |
| HIGD2A | 1.992 |
| FABP7 | 1.915 |
| SLC39A6 | 1.879 |
| HMGB3 | 1.794 |
| CAST | 1.68 |
| TRAPPC1 | 1.623 |
| SRSF4 | 1.584 |
| PABPN1 | 1.582 |
| KHDC4 | 1.541 |
| TRMT112 | 0.657 |
| XAB2 | 0.653 |
| CENPO | 0.646 |
| CIAO2B | 0.644 |
| SNX15 | 0.64 |
| FADS2 | 0.623 |
| RPP38 | 0.622 |
| INPP4A | 0.617 |
| TRIM26 | 0.608 |
| BHMT | 0.567 |
| CCZ1 | 0.54 |
| TRPC4AP | 0.518 |
| TMEM38B | 0.511 |
| NRM | 0.51 |
| TPBG | 0.504 |
| PLCD3 | 0.489 |
| KDM4D | 0.458 |
| NCOA5 | 0.452 |
| EPHA7 | 0.451 |
| SCD | 0.371 |
| WLS | 0.369 |
| ALDH1A1 | 0.309 |
| PUS3 | 0.294 |
| RCL1 | 0.281 |
| GPAA1 | 0.271 |
| FGD2 | 0.136 |
| ALB | 0.097 |

| APOE 2/- vs 3/- |  |
| --- | --- |
| Protein | Fold Change |
| TPPP3 | 1.933 |
| ELAVL4 | 1.831 |
| HMGN1 | 1.743 |
| SH3BGRL | 1.728 |
| FABP7 | 1.66 |
| HMGB3 | 1.632 |
| MAP1B | 1.529 |
| ANXA1 | 0.643 |
| LMNA | 0.624 |
| RPL22 | 0.607 |
| RPP38 | 0.586 |
| KRT18 | 0.58 |
| PUS3 | 0.557 |
| TPD52L1 | 0.469 |
| TPBG | 0.413 |
| SCD | 0.376 |
| WLS | 0.309 |
| ALDH1A1 | 0.292 |
| FGD2 | 0.128 |
| ALB | 0.108 |

CRISPR-edited line APOE 2/- showed 6 DEPs when compared to the parent line, of which 3 were upregulated and 3 were downregulated. CRISPR-edited line APOE 4/- showed 2 DEPs of which 1 was upregulated and 1 was downregulated (Table 2).

### Nucleolar morphology

Several proteins involved in the transcription and processing of rRNA and the biogenesis of ribosomes were differently upregulated upon removal of *APOE4*.

Nucleolus assembles around ribosomal DNA (rDNA) that is being actively transcribed and is an integral part of ribosome biogenesis. Hence, we studied the nucleolar morphology in the parental and isogenic APOE lines. Immunocytochemistry of nucleolar protein Fibrillarin was performed to visualize nucleolar morphology in APOE 3/4 and APOE 3/3 NPCs. The cells were evaluated for the one-nucleolus phenotype, similar to a previous study by [Barnes et al](https://drive.google.com/open?id=1mp5ZcJQpM4Y7V6WePFHLm4u3ZEHFsegI).

[Images and tables](https://docs.google.com/presentation/u/1/d/1o7yHkTdrN90l9OQPPNFxmYu79ELBn1f-/edit)

## DISCUSSION

Multiple studies over the years have shown that APOE is involved in several processes critical for neurodevelopment, including neurogenesis, neuronal migration, and synaptogenesis ([Hack et al., 2007](https://pubmed.ncbi.nlm.nih.gov/17913789/); [Levi et al., 2003](https://www.sciencedirect.com/science/article/abs/pii/S0969996103000457?casa_token=UjrboPUdWCEAAAAA:TRmiDI7NntcWIwPD9VF3FW6VepcR2xa28j_w5yKXk0zZaPo8p5ZOuQuRL-uMs7H4ruO55GRj7E0#preview-section-abstract); [Li et al., 2009](https://www.sciencedirect.com/science/article/pii/S1934590909005268); and thus is essential for the proper development of the brain ([Dean, D.C. et al., 2014](https://pubmed.ncbi.nlm.nih.gov/24276092/); [Yang et al., 2011](https://journals.biologists.com/dev/article/138/20/4351/44686/ApoE-is-required-for-maintenance-of-the-dentate)).

Our analysis shows that APOE plays a role in ribosomal biogenesis, the process by which ribosomes, the cellular machinery responsible for protein synthesis, are produced. We report six DEPs enriched for ribosomal biogenesis and rRNA processing in the *APOE 3/-* compared to the *APOE 3/4* line. Ribosomal biogenesis is a highly complex and energy-demanding process that involves the coordinated assembly of multiple ribosomal RNA (rRNA) and protein components. The process takes place in the nucleolus, a specialized sub-compartment of the nucleus, and requires the involvement of various chaperones, enzymes, and regulatory factors ([Wang et al., 2015](https://www.ncbi.nlm.nih.gov/pmc/articles/PMC4710177/)).

CRISPR-mediated removal of *APOE4* showed differential expression of Nol11. Defective ribosome biogenesis resulting from siRNA depletion of ribosome biogenesis factor NOL11 correlates with changes in nucleolar number in human cells ([Freed et al.,2012](https://drive.google.com/open?id=1TJz9-PYEHgHB3NDolpDlzofUtnt0GEr1)). Nol11 knockdown results in the activation of the nucleolar stress response and craniofacial defects in Xenopus models ([Griffin et al., 2015](https://journals.plos.org/plosgenetics/article?id=10.1371/journal.pgen.1005018)).

CRISPR-mediated removal of *APOE4* also showed differential expression of RCL1. Human RCL1 is highly expressed in astrocytes of the human cerebral cortex ([Amin et al., 2018](https://www.nature.com/articles/mp201749#Abs1)). RCL1 copy number variations show varying neurodevelopmental or psychiatric phenotypes. Dosage variation of RCL1 contributes to a range of neurological and clinical phenotypes ([Brownstein et al.,2021](https://drive.google.com/file/d/12zv7ZfZcEbi5ptU5uclyLFeMqTV4ifhl/view?usp=sharing)).

Additionally, we report the differential expression of four other proteins involved in ribosomal biogenesis that are differentially expressed upon CRISPR-mediated removal of APOE4, such as IPO4, MRM1, RPP38, and TRMT112.

In summary, while further research is needed to fully understand how APOE interacts with the proteins mediating ribosomal biogenesis, this paper highlights that the different isoforms of APOE show differential expression of the proteins involved in this process. Impaired ribosomal biogenesis results in a decrease in protein synthesis capacity and an increase in susceptibility to cellular stress ([Wang et al., 2015](https://drive.google.com/file/d/12zv7ZfZcEbi5ptU5uclyLFeMqTV4ifhl/view?usp=sharing)). Dysfunction in ribosomal biogenesis has been linked to various diseases, including neurodegeneration ([Dönmez-Altuntaş et al., 2005](https://pubmed.ncbi.nlm.nih.gov/16110230/)). Understanding the role of APOE in this process may lead to new insights into the pathogenesis of these diseases.

In neuron..

---

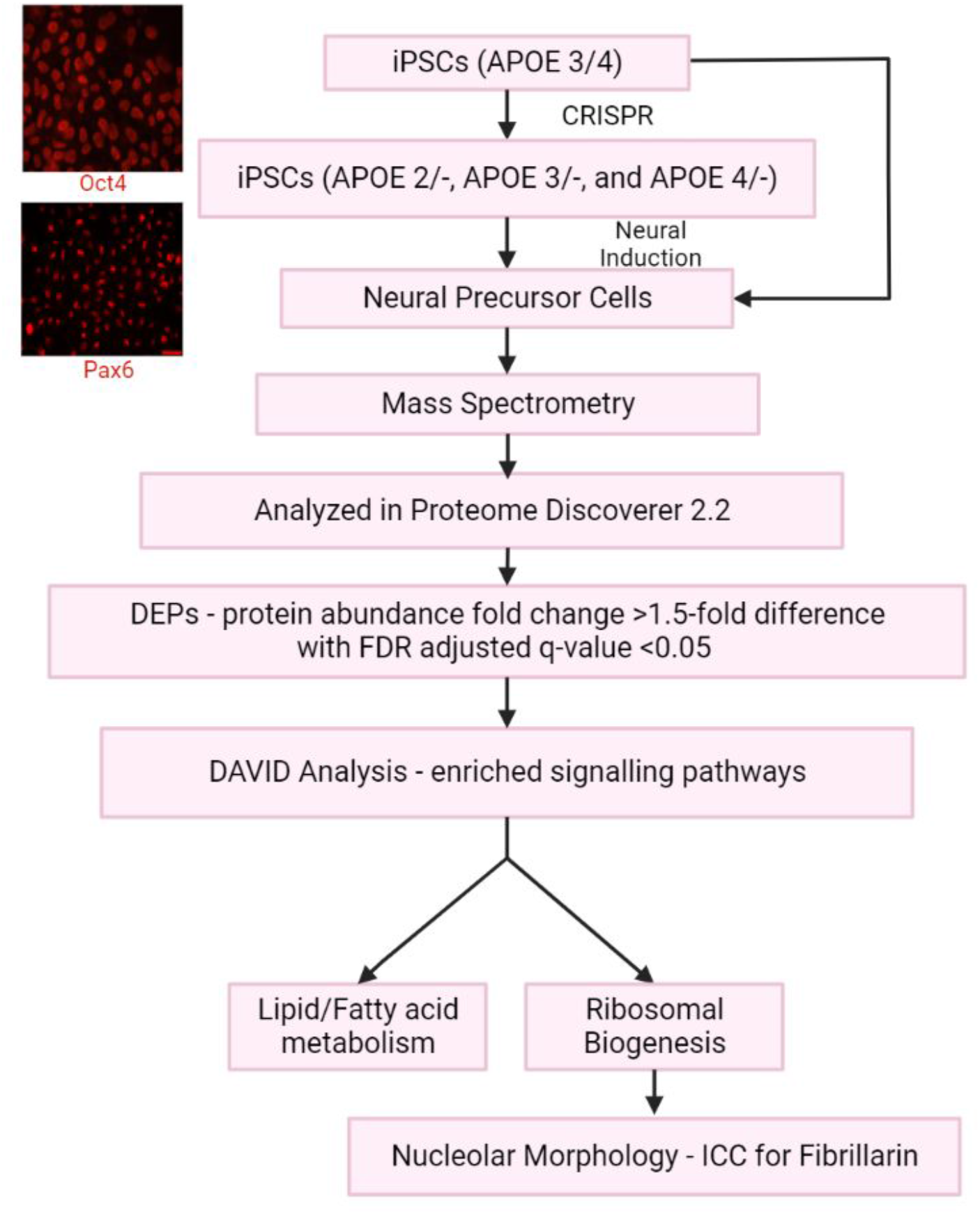

**Fig 1:**
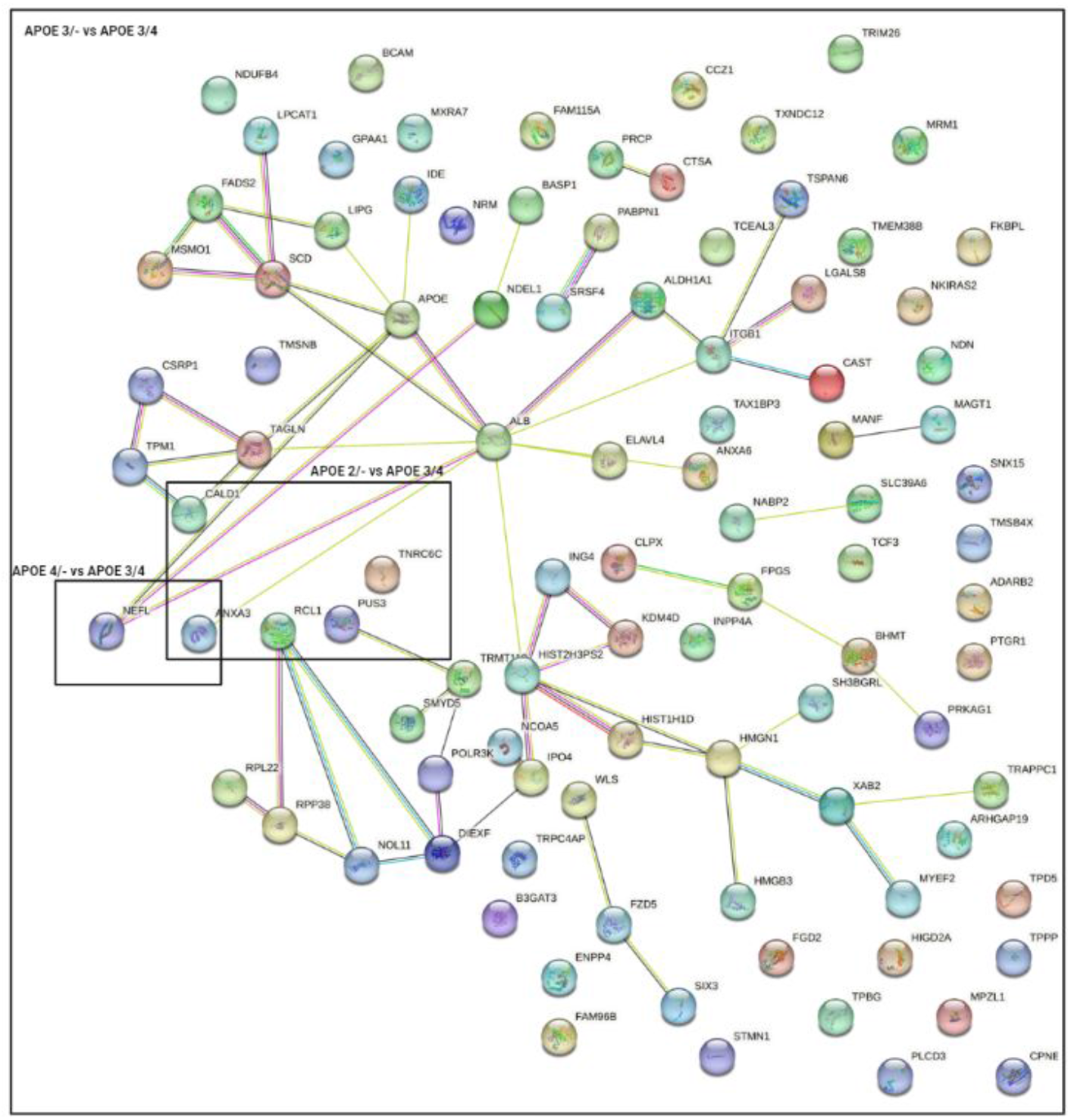
Interactome of DEPs in APOE2/-, APOE3/-, and APOE4/- compared to the parental line

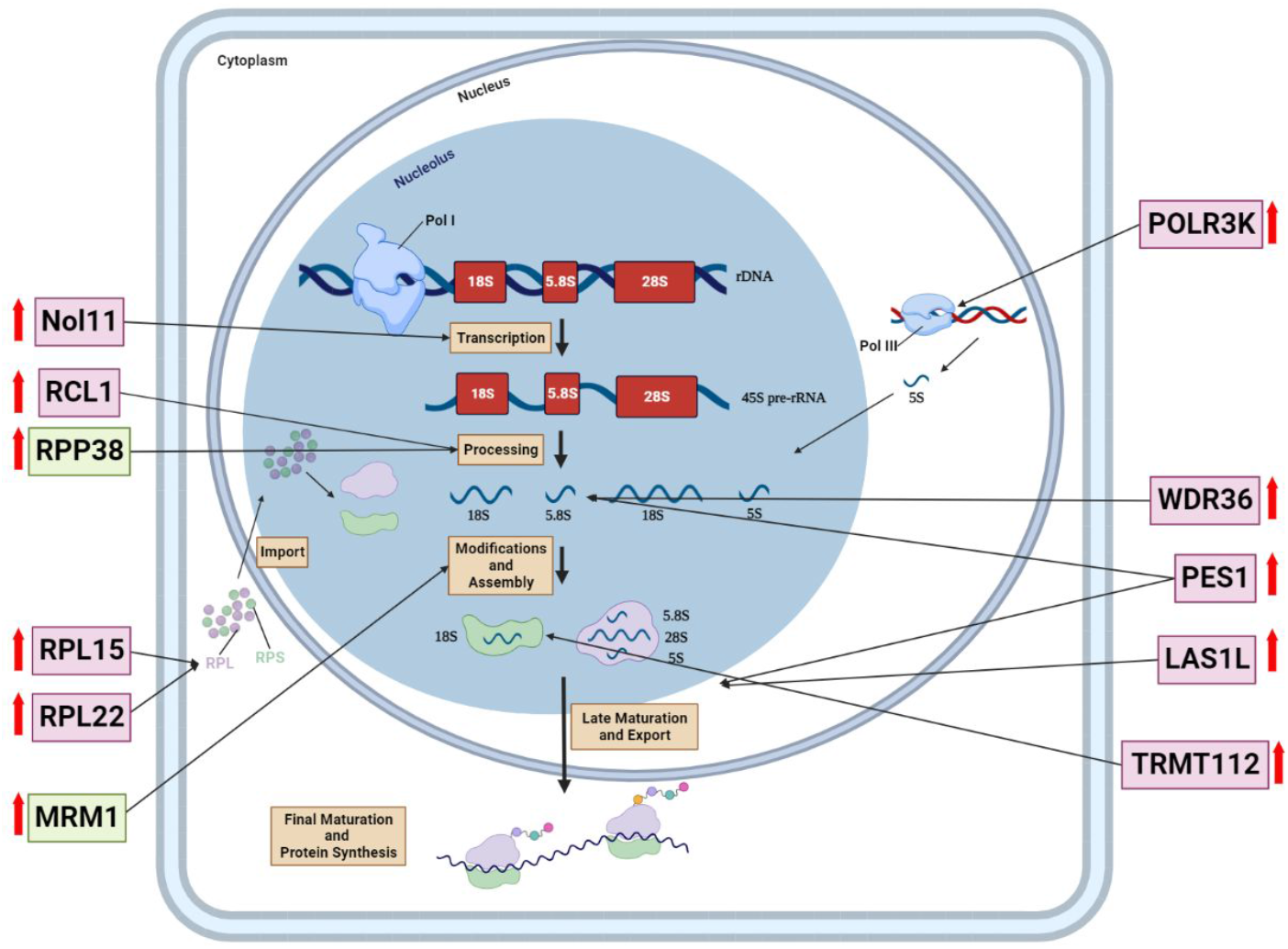

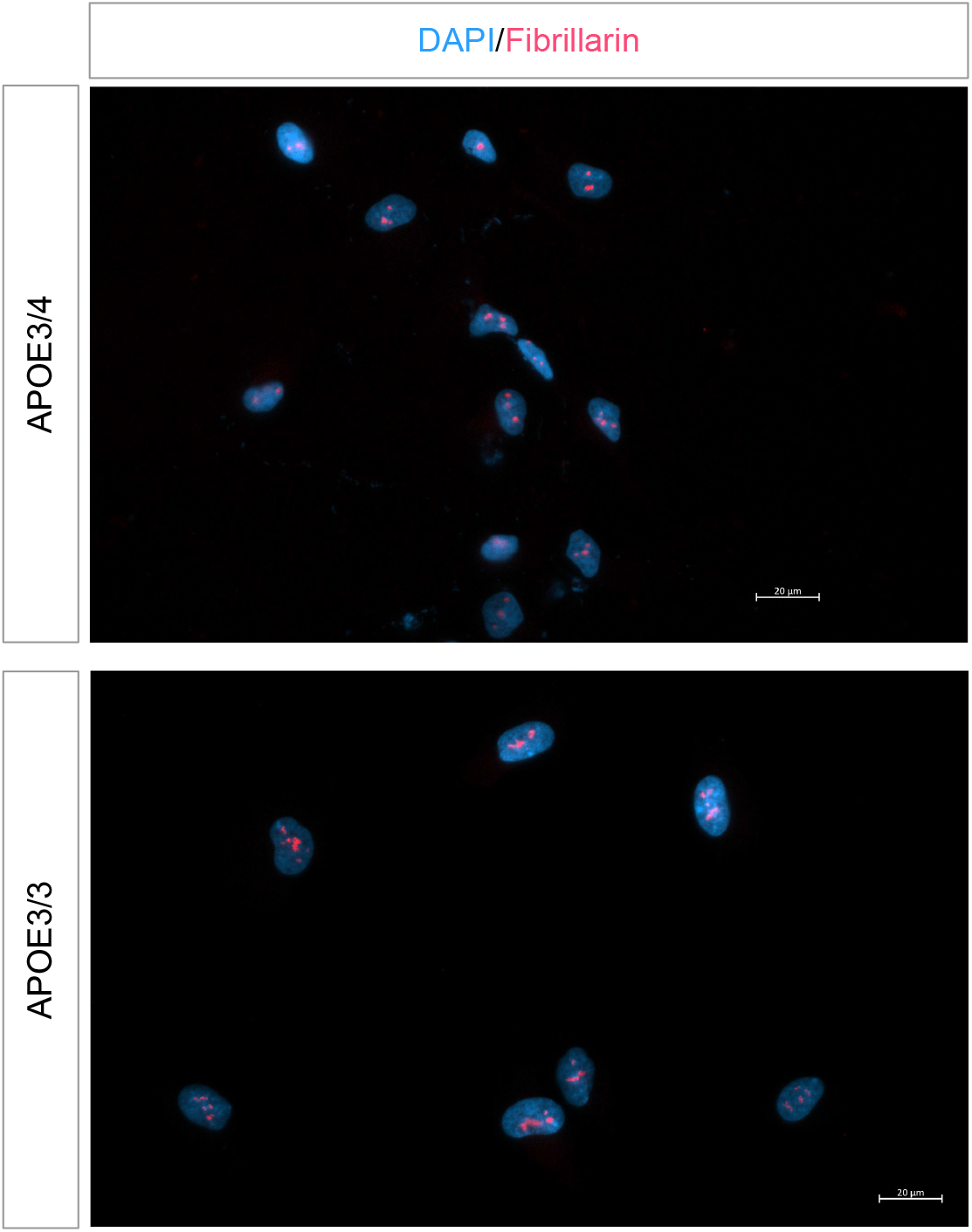

